# Unravelling the Role of *Candida albicans* Prn1 in the Oxidative Stress Response Through a Proteomic Approach

**DOI:** 10.1101/2023.11.07.566035

**Authors:** Víctor Arribas, Lucía Monteoliva, María Luisa Hernáez, Concha Gil, Gloria Molero

## Abstract

*Candida albicans* Prn1 is a protein that shares similarities with mammalian Pirin but with an unknown function in the yeast. Orthologues of Prn1 have been identified in other pathogenic fungi but not in *Saccharomyces cerevisiae*, suggesting a relationship with pathogenesis. Prn1 increase in abundance after H_2_O_2_ treatment has been shown previously, thus, in the present work, *C. albicans prn1Δ* mutant and the corresponding wild-type strain SN250 have been treated with H_2_O_2_ and their response was studied by quantitative differential proteomics. These assays indicated a lower increase of proteins with oxidoreductase activity after treatment in the *prn1Δ* strain compared to the wild type, as well as an increase in proteasome-activating proteins and a decrease in translation-involved proteins. Accordingly, Prn1 absence, under H_2_O_2_ treatment, led to a lower survival rate and a higher percentage of apoptosis, together with higher reactive oxygen species levels and higher proteasome activity. Besides, remarkable differences in the abundance of some transcription factors were observed between the two strains. Mnl1, involved in Prn1 expression, Bas1, Tiff33, and orf19.1150 presented an inverse pattern of expression under H_2_O_2_ treatment respect to Nrg1, a Mnl1 antagonist. Interestingly, orf19.4850, a protein orthologue to *S. cerevisiae* Cub1, has shown to be involved in the response to H_2_O_2_ presenting a conserved proteasome function. Under basal conditions, the proteomics results indicate a possible involvement of Prn1 in mitochondrial oxidative stress detoxication. Our experiments confirm Prn1 as a relevant actor in the oxidative response.

**Importance:** *Candida albicans* is a human opportunistic pathogen included in the WHO fungal priority pathogens list. The increase in resistant strains necessitates the discovery of new targets for antifungal therapies. Our research sheds light on the important role of the previously uncharacterized *C. albicans* protein Prn1 during the oxidative stress response. Study of the proteome remodelling under oxidative stress unveils the role of Prn1 in the decreased reactive oxygen species levels and the consequences, such as death by apoptosis and necrosis or cell growth delay. A proteomics approach allowed the identification of several proteins potentially involved in Prn1 activity, such as oxidoreductases and transcription factors. The lack of Prn1 orthologues in *Saccharomyces cerevisiae* but the presence in other *Candida* and *Aspergillus* species implicates this protein in pathogenesis and suggests that it may serve as a candidate for new drug targets.

## Introduction

*Candida albicans* is a dimorphic fungus which is part of the commensal human microbiota. However, under certain conditions, this opportunistic pathogen causes systemic infections, especially in immunocompromised patients (1, 2). Invasive candidiasis is one of the main nosocomial diseases, particularly among patients in intensive care units, making it a major public health concern. As a result, the World Health Organization (WHO) has included this pathogen in their priority list of fungal pathogens (3–5).

During invasive candidiasis, *C. albicans* interacts with phagocytes, mainly macrophages. Macrophages produce oxidant molecules, such as hydrogen peroxide (H_2_O_2_) and nitric oxide (NO) in response to the infection. Successful *C. albicans* infection depends, to a significant degree, on the resistance to these oxidative agents (6–8). To detoxify reactive oxygen species (ROS), three main mechanisms are activated by the Cap1 and Hog1 signalling pathways in *C. albicans*, involving the catalase, glutathione, and thioredoxin systems (9, 10). Experiments from our group have shown that *C. albicans* exhibits important proteome remodelling to counteract the oxidative attack by macrophages. These changes include enrichment of proteins with oxidoreductase and superoxide dismutase activities, which counteract the high ROS levels in the cell. Yeast cells also experience an increase in the metabolism of amino acids and nucleotides, in contrast to a reduction in glycolysis and in translation (11). Ultimately, these changes lead to 30% of *C. albicans* cells to undergo programmed cell death by apoptosis (8). The treatment of *C. albicans* with H_2_O_2_ provides an excellent strategy to reproduce the oxidative stress induced by phagocytes. This agent was used in a recent quantitative proteomic analysis allowing the identification of new proteins that are likely involved in the detoxification process, such as Prn1 or Oye32 (12).

Prn1 is a protein that shares similarities with mammalian Pirin but with an unknown function in *C. albicans*. Pirin expression correlates with the activation of antioxidant transcription factor Nrf2 and the subsequent expression of related proteins, such as NAD(P)H oxidoreductase 1 (NQO1) (13, 14). Pirin also contains a cupin-activating domain that enables binding to metal ions, such as iron or copper, changing its conformation from the inactive form (Fe^2+^ binding) to the active form (Fe^3+^ binding) (15). This activation is crucial to modulate the response to different stresses, such as oxidative stress or cell death by apoptosis (16). A similar protein, PirA, has been identified in a prokaryotic microorganism, *Streptomyces*, as a negative regulator of the mitochondrial beta-oxidation pathway involved in the reduction of oxidative stress (17). Thus, Prn1 may be related to the oxidative stress response. In *C. albicans, PRN1* has three homologues: *PRN2, PRN3*, and *PRN4* being the last one the most similar to *PRN1*. On the other hand, while some uncharacterized orthologues are predicted in other *Candida* species and *Aspergillus nidulans*, no orthologue is found in *S. cerevisiae*, suggesting that Prn1 is important for pathogenesis.

To shed light on the role of Prn1 in the oxidative stress response, we used a data-dependent acquisition (DDA) quantitative proteomics approach after H_2_O_2_ treatment in a null *prn1Δ* mutant and the corresponding wild-type strain (SN250) (18). Prn1 absence leads to a smaller increase in proteins related to the oxidative response and a greater decrease in translation-related proteins. This would probably be the cause of the observed increased in ROS levels and the increased level of apoptosis in the deleted strain with respect to the wild-type. To counteract the loss of Prn1, *C. albicans* seems to increase other molecular mechanisms that include greater proteasome activity. In addition to its importance during oxidative stress, under basal conditions, Prn1 may also be implicated in mitochondrial cell redox. In addition, transcription factor Mnl1 exhibits an opposite behaviour between wild-type and the deleted strain, being detected under oxidative stress in *prn1Δ* but not in the wild-type strain. We have also identified the previously undescribed *C. albicans* protein corresponding to orf19.4850, a gene orthologue to *S. cerevisiae CUB1*, which had increased abundance only in the wild-type, in which Prn1 is expressed under oxidative stress.

## Results

### Impact of *PRN1* deletion on cell survival and recovery under oxidative stress

Cell survival during oxidative stress was quantified by propidium iodide (PI) staining after 200 min in the presence of 10 mM H_2_O_2_ for the *prn1Δ* and SN250 strains. Flow cytometric analyses showed that up to 45% of cells in the *prn1Δ* strain were dead cells after treatment compared with 35% for the SN250 strain (Fig. 1a). This increased percentage of dead cells correlates with the delay in cell growth of nascent cultures of both strains after exposure to H_2_O_2_ in a dose-dependent manner. This delay was more pronounced for the *prn1Δ* strain (Fig. 1b). These experiments show the importance of Prn1 for the viability and recovery of *C. albicans* after H_2_O_2_ treatment.

**FIG 1.**
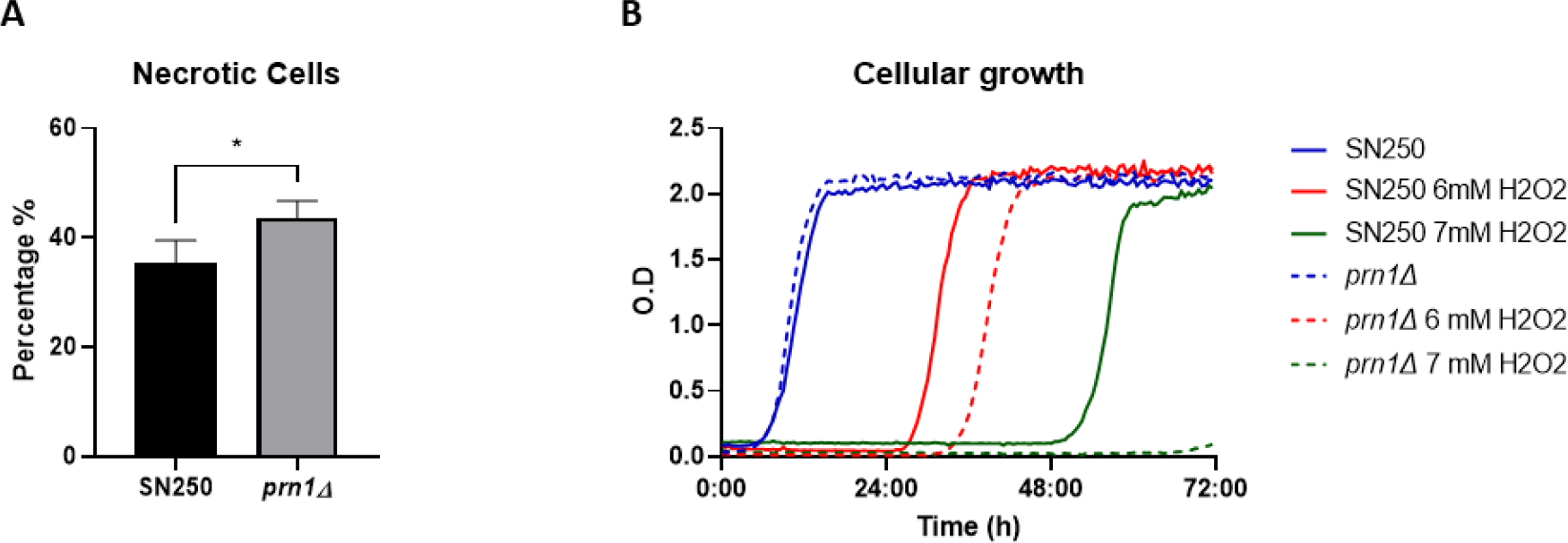
SN250 and *prn1Δ* strain viability and growth in the presence of H_2_O_2_. (a) Percentage of propidium iodide (PI) positive necrotic cells measured by flow cytometry after 200 min in the presence of 10 mM H_2_O_2_. Results represent the average of three biological replicates. Error bars indicate standard deviation. * p <0.05, unpaired t-test. (b) Growth curves of both strains in the presence of 6 mM and 7 mM H_2_O_2_. The graph presents the most representative curve of three biological replicates.

### Proteomic response of SN250 and *prn1Δ* strains to H2O2 treatment

To unmask the role of Prn1 in the oxidative stress response, we performed a quantitative proteomics assay (DDA-MS) of the SN250 and *prn1Δ* strains after 200 min of H_2_O_2_ treatment and compared it to the control condition. The proteomics analyses allowed the quantification of approximately 1,800 *C. albicans* proteins on each strain. Statistical analysis of the relative quantification (q-value <0.05) revealed that 176 and 183 proteins significantly changed their abundance in response to the treatment of SN250 and *prn1Δ*, respectively (Fig. 2a and Table S1). The volcano plots in Figure 2b represent the different changes in protein abundance between the two strains. As shown in the Venn diagram (Fig. 2c), only 98 proteins were common to both comparisons, highlighting the importance of Prn1 in the oxidative response (Table S2).

**FIG 2.**
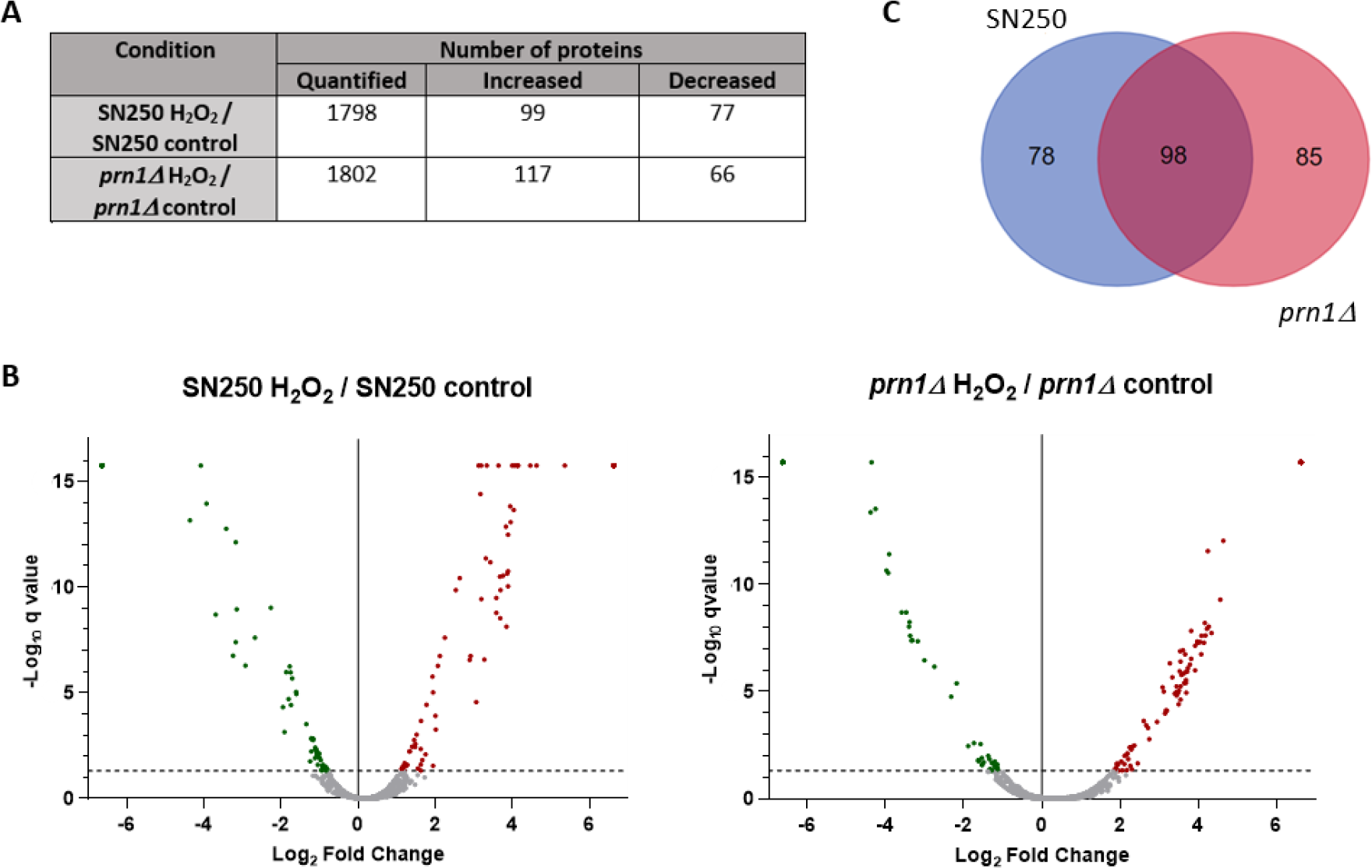
Quantitative proteomics assay (DDA-MS) of SN250 and *prn1Δ* strains in response to 200 min of 10 mM H_2_O_2_ treatment. (a) Number of proteins quantified and that showed significant differences in abundance between the two culture conditions for each strain. (b) Volcano plots representing proteins with significant changes in abundance. Significant changes in the protein abundance (-log10 q-value >1.3) after treatment are presented in red for increased or green for decreased. (c) Venn diagram showing proteins with significant changes in abundance in each strain or both.

Analysis of the 10 proteins with the greatest increase in the relative abundance for each strain only revealed three proteins in common between them: Cip1, Ach1, and Pim1 (Table 1). In the SN250 strain, we observed previously described proteins (e.g., Prn1 or Oye22) (12), as well as other proteins that did not significantly increase in the *prn1Δ* strain (e.g., Psa2 and Alt1). For the *prn1Δ* strain, we detected the Qcr9 and Nuc2 proteins implicated in ubiquinone/ubiquinol redox. Qcr9 did not significantly increase in abundance in the wild-type strain, as also occurs with orf19.7310. Particularly interesting was orf19.6035 which abundance significantly increased in both strains but its function remains unknown.

**TABLE 1.** Top 10 proteins with the greatest increase in relative abundance in after treatment in the SN250 or *prn1Δ* strain compared with the control condition and the other strain.

| Strain | Protein | Description | SN250 H<sub>2</sub>O<sub>2</sub> /<br>SN250 control<br>ratio log<sub>2</sub> | *prn1Δ* H<sub>2</sub>O<sub>2</sub> /<br>*prn1Δ* control<br>ratio log<sub>2</sub> |
| --- | --- | --- | --- | --- |
| **SN250** | Psa2 | Mannose-1-phosphate guanylttransferase | 5.36 | NS |
|  | Icl1 | Isocitrate lyase; glyoxylate cycle enzyme | 4.63 | 3.82 |
|  | Prn1 | Protein with similarity to pirins | 4.16 | ND |
|  | Alt1 | Putative alanine transaminase | 4.13 | NS |
|  | Oye22 | Putative NADPH dehydrogenase | 4.06 | 3.28 |
|  | Acc1 | Putative acetyl-coenzyme-A carboxylases | 4.04 | 3.57 |
|  | orf19.3477 | Putative pseudouridine synthase | 4 | 3.54 |
| **SN250**<br>**&**<br>***prn1Δ*** | Cip1 | Possible oxidoreductase | 4.47 | 4.65 |
|  | Ahc1 | Ortholog(s) histone acetyltransferase activity | 4.13 | 4.23 |
|  | Pim1 | ATP-dependent Lon protease | 3.96 | 4.17 |
| ***prn1Δ*** | Mal2 | Alpha-glucosidase | 4.57 | + |
|  | Lys21 | Homocitrate synthase | 4.34 | 3.07 |
|  | orf19.6035 | Protein of unknown function | 4.27 | 3.89 |
|  | Hsp21 | Small heat shock protein | 4.25 | 3.19 |
|  | Qcr9 | Putative ubiquinol cytochrome c reductase | 4.19 | NS |
|  | Nuc2 | Putative NADH-ubiquinone oxidoreductase | 4.15 | + |
|  | orf19.7310 | Role in directing meiotic recombination | 4.09 | NS |

“+” indicates proteins detected only in the treated condition, “NS” indicates non-significant change in abundance and “ND” indicates non-detected proteins.

Gene Ontology (GO) enrichment for the biological process and function of all proteins with significantly increased abundance in each strain (Table S1) showed a high increase in proteins with the oxidoreductase activity GO term for both strains (Fig. 3a). Many of these proteins were different for each strain, and the number was higher for SN250 (Table S3). We also observed enrichment of protein catabolic process GO term proteins for both strains in the presence of H_2_O_2_ (Fig. 3b). Increased proteins in the *prn1Δ* strain were mainly related to the regulatory subunit of the proteasome (Phb2, Pr26, Rpn3, and Rpt4), whereas these proteins in the SN250 strain were chaperones and ubiquitin-binding proteins involved in protein transport from Golgi to vacuole (Bzz1, orf19.4430, Vps4, and Mdj1). The *prn1Δ* strain also presented a greater decrease in the abundance of proteins associated with translation than the treated SN250 strain (Fig. 3c). Both strains presented with a decrease in nucleotide metabolic process GO term proteins, which are related to purine and pyrimidine biosynthesis, and this effect was higher in the *prn1Δ* strain (Fig. S1). To emphasize the differential response of each strain to oxidative stress, GO term analysis was carried out for the proteins that exclusively increased in abundance for each strain (Fig. 2c). The 78 proteins that significantly increased in abundance only in the SN250 strain were significantly enriched in oxidoreductase activity proteins, suggesting a higher oxidative stress response when Prn1 is present. On the other hand, GO term enrichment of the 85 proteins that significantly increased in abundance only in the *prn1Δ* strain was related to proteasome-activating activity, which could indicate higher proteasome activity (Fig. S2).

**FIG 3.**
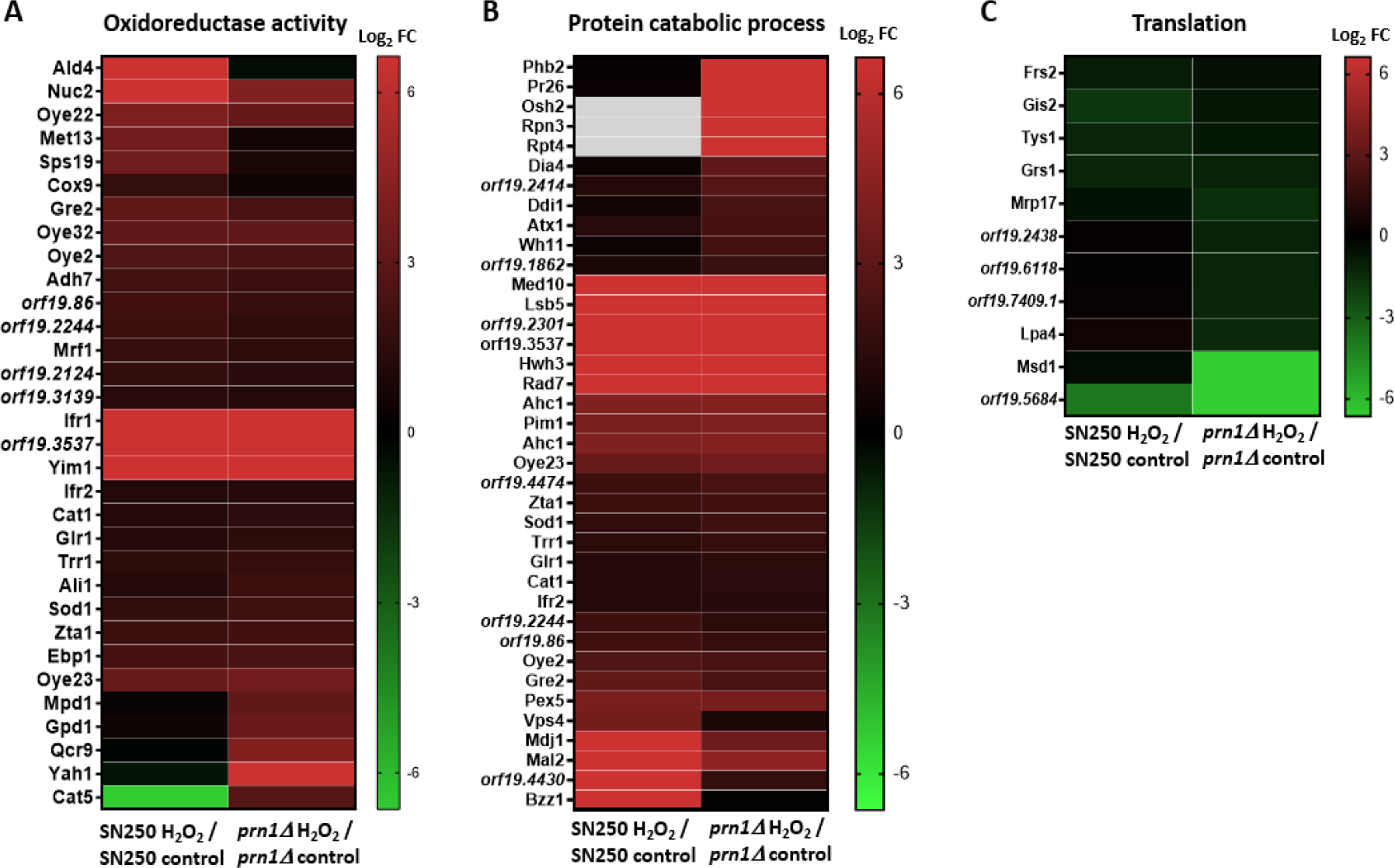
Hierarchical clustering heat maps of the proteins that changed in abundance after 200 min of 10 mM H_2_O_2_ treatment in each strain with respect to the control condition grouped by GO term. (a) Oxidoreductase, (b) protein catabolic process, (c) translation. Grey gaps indicate proteins not detected in that strain.

The proteins that increased in abundance in the SN250 and *prn1Δ* strains under oxidative treatment (Table S1) were grouped into 13 and 11 predicted protein network clusters, respectively, using STRING software. In the SN250 strain, we found three clusters related to oxidoreductase function. Other clusters were implicated in pre-ribosome and ribosome biogenesis, dehydrogenase complex of the respirasome, and amino acid biosynthesis (Fig. 4a). For the *prn1Δ* strain, we also found protein clusters involved in the oxidative stress response and ribosome biosynthesis, but also in the proteasome regulatory subcomplex, heat shock response, mitochondrial oxidoreductase complex, and Rho GTPase regulation (Fig. 4b).

**FIG 4.**
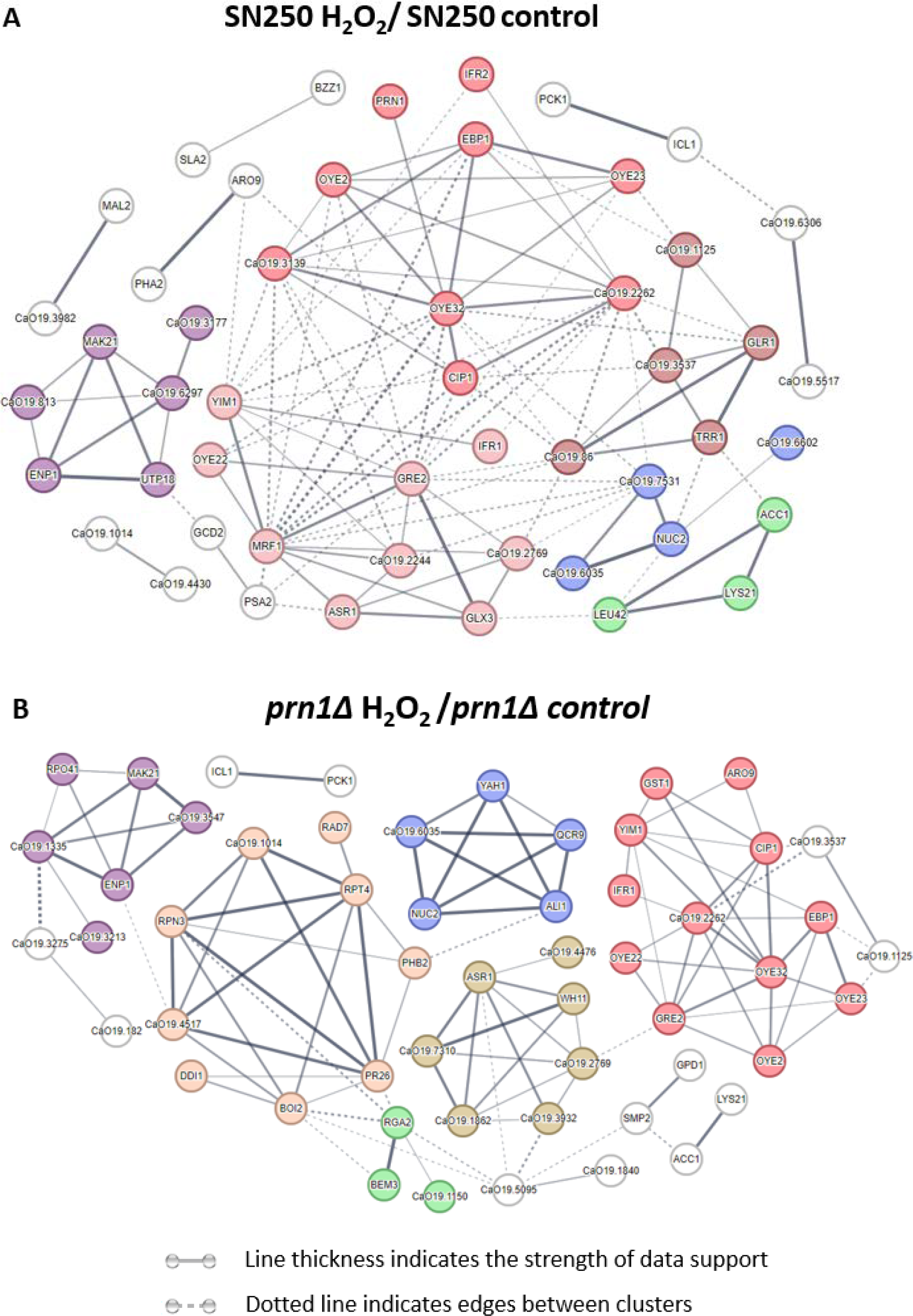
Predicted networks of proteins with increased abundance after H_2_O_2_ treatment in the SN250 and *prn1Δ* strains using STRING software. Line thickness indicates the strength of supporting data. Dotted lines indicate edges between clusters. (a) Clusters: oxidoreductase function (red, dark red, and light red), preribosome and ribosome biogenesis (purple), dehydrogenase complex of the respirasome (blue), and amino acid biosynthesis (green). (b) Clusters: oxidative stress response (red), proteasome regulatory subcomplex (orange), heat shock response (gold), ribosome biosynthesis (purple), mitochondrial oxidoreductase complex (blue) and Rho GTPase regulation (green).

Proteins only detected in one condition but not in the other for each strain are presented in Table S4. Among them, we selected those that did not show any significant change or were not detected in the other strain (Table 2). In the case of the SN250 strain, we highlight riboflavin and carnitine antioxidant biosynthesis-related proteins (Rib2 and Ald4). Interestingly, orf19.4850, a gene orthologue to *S. cerevisiae CUB1* linked to DNA repair and proteasome function (19), encodes a protein that was only detected in the wild-type strain with H_2_O_2_ treatment. For the *prn1Δ* strain, we highlight proteins related to catabolism (Osh2, Phb2, Pr26, Rpn3, and Rpt4) and two transcription factors, Tif33 and orf19.1150, that were only detected under oxidative stress. Conversely, among the proteins that were not detected after oxidative treatment in the *prn1Δ* strain, we found proteins related to ribosome biogenesis (Afg2, Has1, Lig1, and Ubp12) and the transcription factor/repressor Nrg1.

**TABLE 2.** Proteins detected in one condition but not in the other and without significant change in abundance or not detected in the other strain. Protein descriptions and quantification of relative abundance are found in Table S4.

| Strain | Detected in treated condition | Detected in basal condition |
| --- | --- | --- |
| SN250 H<sub>2</sub>O<sub>2</sub> / SN250 control | Ald4, Bzz1, orf19.4430, orf19.4850, Rib2 | Hrr25, Mak11, Mcm3, orf19.1272, Ynd1 |
| *prn1Δ* H<sub>2</sub>O<sub>2</sub> / *prn1Δ* control | Fmp28, Nit2, orf19.1150, orf19.2051, orf19.2452, Osh2, Phb2, Pr26, Rpn3, Rpt4, Rts1, Smp2, Snu66, Tad2, Tif33, Uso1, Yah1 | Afg2, Has1, Lig1, Msd1, Nrg1, orf19.5987, orf19.6445, Pnc1, Rpa43, Sec24, Trp1, Ubp12 |

The most dramatic differences in protein abundance from Table S4 are those with an opposite change between both strains after H_2_O_2_ treatment (Table 3). Among these proteins, we observed two transcription factors (Mnl1 and Bas1), Mfg1 biofilm growth regulator, and two ubiquinone biosynthesis-related proteins (Cat5 and Fmp53).

**TABLE 3.** Proteins that presented opposite changes in abundance between both strains after H_2_O_2_ treatment.

| Protein | Description | SN250 H<sub>2</sub>O<sub>2</sub> / SN250 control Ratio log<sub>2</sub> | *prn1Δ* H<sub>2</sub>O<sub>2</sub> / *prn1Δ* control Ratio log<sub>2</sub> |
| --- | --- | --- | --- |
| Bas1 | Putative helix-loop-helix (HLH) transcription factor; role in filamentous growth | - | 3.61 |
| Cat5 | 2-octoprenyl-3-methyl-6-methoxy-1,4-benzoquinone hydroxylase activity; ubiquinone biosynthetic process | - | 2.75 |
| Dad4 | Subunit of the Dam1 (DASH) complex; chromosome segregation | -3.15 | 4.06 |
| Fmp53 | Ubiquinone-6 biosynthetic process | 3.69 | - |
| Ftr2 | High-affinity iron permease | -2.67 | 2.11 |
| Mfg1 | Regulator of filamentous growth; biofilm formation | - | 3.5 |
| Mnl1 | Transcription factor; induces transcripts of stress response genes via SLE (STRE-like) elements | - | + |
| orf19.764 | Ortholog(s) negative regulation of TORC1 signalling | - | + |
| Rpo41 | Putative mitochondrial RNA polymerase; repressed in core stress response | - | 3.45 |

“+” indicates proteins only detected in the treated condition, “-” indicates proteins only detected in the control condition.

### Comparative proteomics analysis between the SN250 and *prn1Δ* strains in the presence or absence of oxidative stress

Comparative analysis of the proteome between both strains in the absence or presence of oxidative stress highlights the possible role of Prn1 under both circumstances. Statistical analysis (q-value <0.05) uncovered only 47 proteins with a significant change in abundance between both strains without an oxidative agent and 67 proteins after treatment with H_2_O_2_, showing that the SN250 strain had more proteins with an increased abundance in response to H_2_O_2_ (Fig. 5a and Table S5). Volcano plots show the greater number of proteins with significant changes in abundance after H_2_O_2_ treatment compared with the untreated condition (Fig. 5b), supporting an important role of Prn1 in response to this stress.

**FIG 5.**
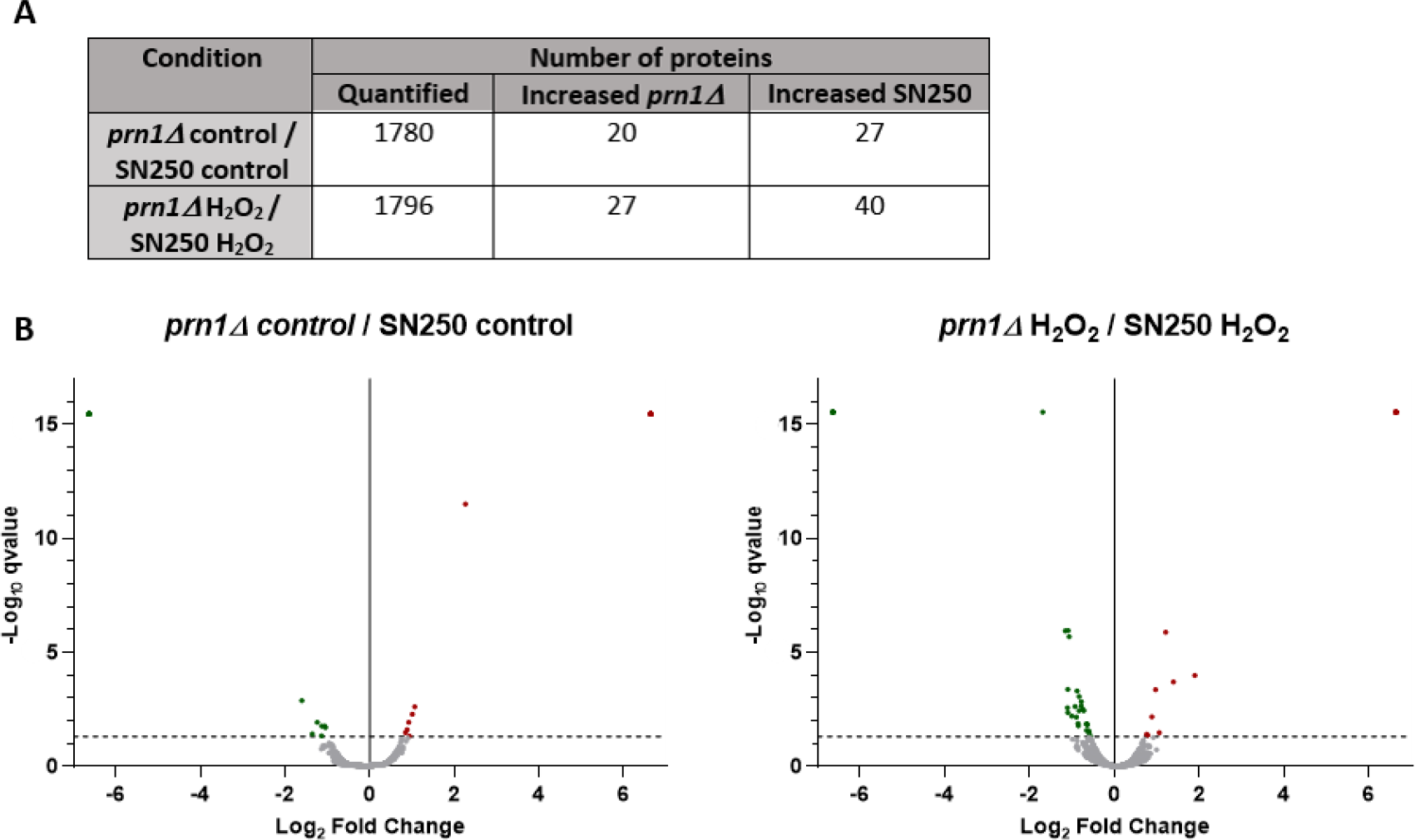
Comparative proteomics analysis between the SN250 and *prn1Δ* strains in the control condition and after 200 min of 10 mM H_2_O_2_ treatment. (a) Number of quantified proteins and those with differences in abundance for each strain. (b) Volcano plots representing proteins with a significantly different abundance between the two strains in each condition.

The mitochondrial component GO term proteins with significant changes in abundance between both strains under control conditions are presented in Figure S3a. The proteins with significantly increased abundance in the *prn1Δ* strain were related to mitochondrial ribosome (Img1 and orf19.2214) and mitochondrial chaperones (Mdj1 and Hsp78), whereas those in the SN250 strain were related to protein import into mitochondria (Fmp28 and Tom6), mitochondrial oxidoreductase/cell redox (Phb2 and Yah1), and mitochondrial respiratory chain assembly (orf19.1336.2). This analysis suggests an important role of Prn1 in mitochondrial redox regulation under basal conditions. In addition, two transcription factors (Mnl1 and orf19.1150) and Swi3, a protein involved in chromatin remodelling, exhibited significant changes.

Under oxidative stress, GO term enrichment analysis of the proteins with a significant change in abundance between both strains indicated an increase in proteasome-activating activity proteins for the *prn1Δ* strain and in translation for the SN250 strain (Fig. S3b), confirming the previously described results.

### Influence of Prn1 deletion on ROS production and proteasome function

The GO term enrichment analysis of proteins that vary in abundance after H_2_O_2_ treatment when Prn1 is present suggested that this protein may play an important role in regulating the detoxification process. To functionally validate this result, we measured intracellular ROS levels using a dihydrorhodamine 123 (DHR123) probe after 200 min of 10 mM H_2_O_2_ treatment. The ROS levels were significantly higher in the Prn1-null mutant relative to the wild-type strain (Fig. 6a). Oxidative stress-mediated ROS accumulation induces apoptotic cell death (12). To evaluate the first signs of apoptosis, both the *prn1Δ* and SN250 strains were treated with 10 mM H_2_O_2_ for 50 min, and phosphatidylserine (PS) externalization, a marker of apoptosis, was measured by flow cytometry. Annexin V staining showed that 34% of treated *prn1Δ* cells presented PS externalization, in contrast to 18% of treated SN250 cells (Fig. 6b). As described previously, GO term enrichment showed an increase in proteasome-activating activity proteins in the *prn1Δ* strain. Therefore, we examined the activity of the proteasome complex by measuring the chymotrypsin-like protease activity after 200 min of 10 mM H_2_O_2_ treatment. Both strains significantly increased the proteasome activity due to the oxidative treatment, with a significantly higher increase for the *prn1Δ* strain (Fig. 6c) in concordance with the results of the proteomics assay.

**FIG 6.**
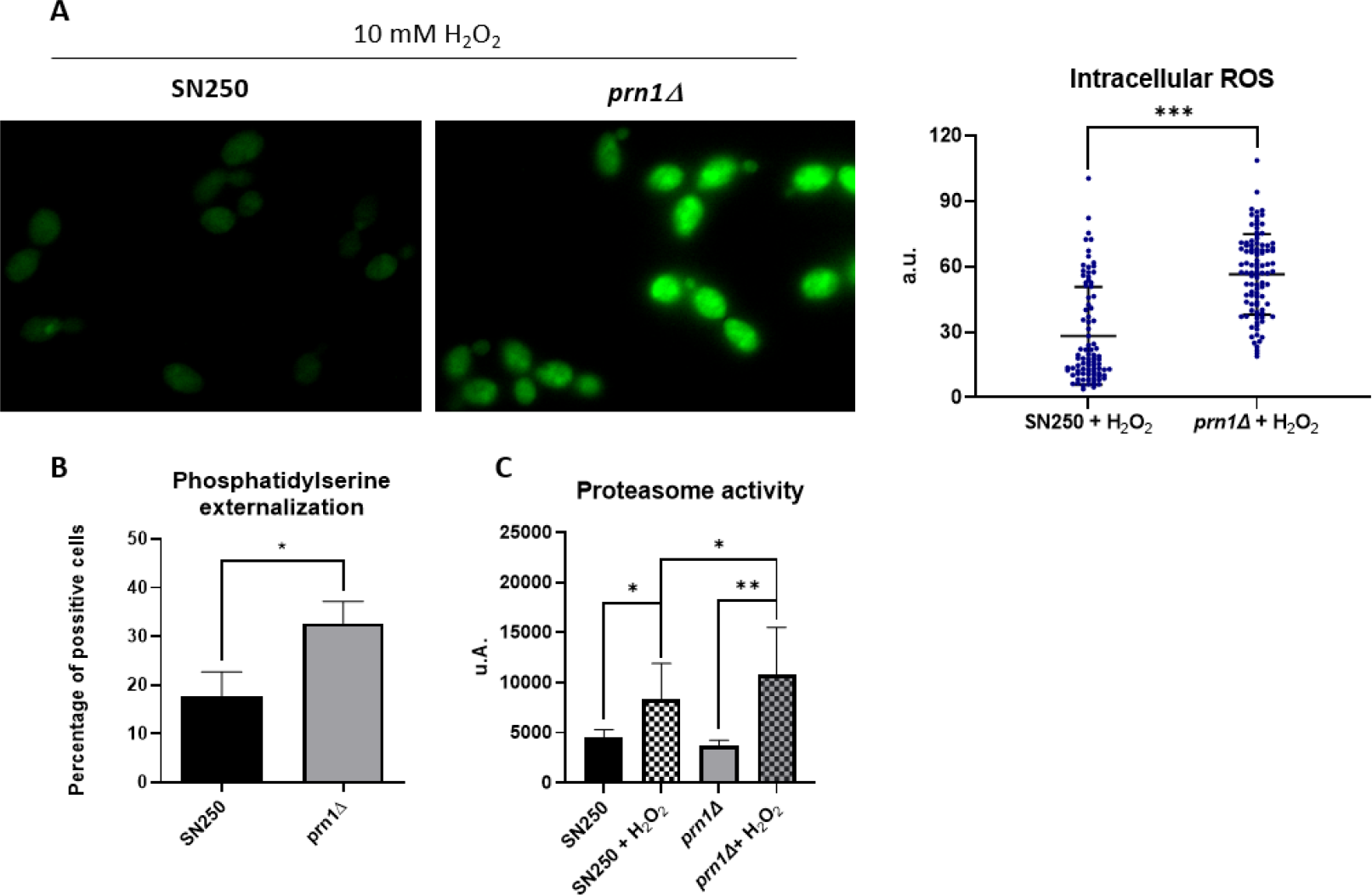
Effect of Prn1 deletion on cell redox homeostasis and apoptosis. (a) Intracellular ROS levels in the SN250 and *prn1Δ* strains after 200 min 10 mM H_2_O_2_ treatment using the dihydrorhodamine 123 (DHR123) probe. (b) Phosphatidylserine externalization levels after 50 min 10 mM H_2_O_2_ treatment using Annexin V staining. (c) Chymotrypsin-like proteasome activity after 200 min 10 mM H_2_O_2_. Results represent the averages of three biological replicates. Error bars indicate standard deviation. ^*^ p <0.05, ^**^ p <0.01, ^***^ p <0.001, unpaired t-test.

### *C. albicans* orf19.4850 mutant response to oxidative stress

The detection of protein corresponding to the uncharacterized *C. albicans* orf19.4850, a gene orthologue of *S. cerevisiae CUB1*, exclusively in the SN250 strain under oxidative stress suggests that its expression could be related to both to oxidative stress and the presence of Prn1. We measured the susceptibility of the *prn1Δ/PRN1* and *cub1Δ/CUB1* mutant strains and the corresponding wild-type *C. albicans* SC5314 to 80 mM H_2_O_2_ in a drop growth assay on YPD medium. The growth of both mutant strains was notably reduced (Fig. 7a), and flow cytometric analyses after 200 min of 10 mM H_2_O_2_ treatment showed that necrotic cells significantly increased for both mutant strains with respect to the wild-type strain (Fig. 7b). The proteasome activity of the *cub1Δ/CUB1* strain after 200 min of 10 mM H_2_O_2_ treatment also increased significantly with respect to its wild-type control strain (Fig. 7c). Thus, the *cub1Δ/CUB1* strain showed a similar behaviour as the *prn1Δ* mutant strain under oxidative stress.

**FIG 7.**
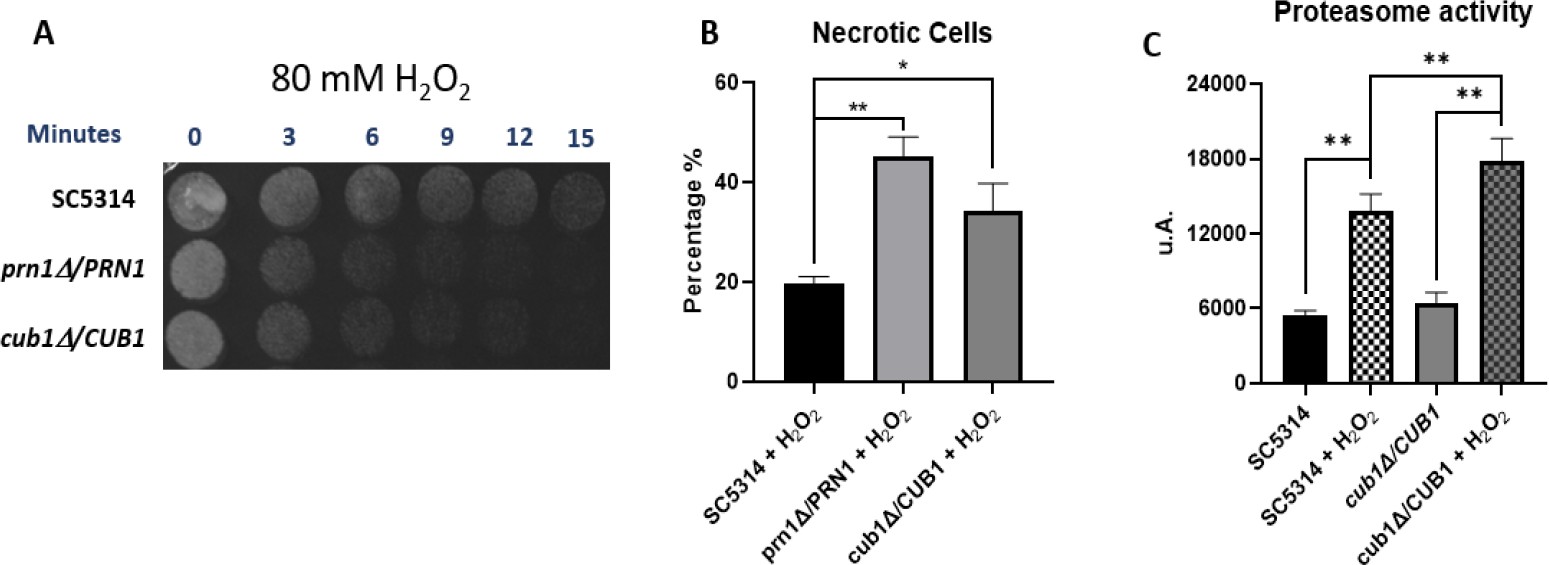
Role of the *C. albicans* Cub1 orthologue protein in the oxidative stress response after H_2_O_2_ exposure. (a) Drop growth assay of *C. albicans* SC5314, *prn1Δ/PRN1, and cub1Δ/CUB1* strains treated with 80 mM H_2_O_2_ for different time intervals. (b) Percentage of PI-positive necrotic cells measured by flow cytometry after 200 min in the presence of 10 mM H_2_O_2_. (c) Chymotrypsin-like proteasome activity after 200 min 10 mM H_2_O_2_ treatment. Results represent the averages of three biological replicates. Error bars indicate standard deviation ^*^ p <0.05, ^**^ p <0.01, unpaired t-test.

## Discussion

### Prn1 has a main role in oxidative response

In a previous quantitative proteomics study in *C. albicans* treated with H_2_O_2_, Prn1 highly increased in abundance after H_2_O_2_ treatment (12). The function of Prn1 in *C. albicans* remains unknown, but the protein presents partial homology with human Pirin, suggesting a possible conserved function (Fig. S4). In mammalian cells, Pirin has a main role in the oxidative stress response through its regulation by the oxidative stress response transcription factor Nrf2 (14). In this study, the *prn1Δ* strain showed an increase in necrotic cells after H_2_O_2_ treatment (Fig. 1a) and a delay in cell growth in response to this agent (Fig. 1b). Oxidative stress in *C. albicans* promotes activation of the Hog1 signalling pathway (9), leading to a transient G1 cell cycle arrest (20). Thus, the observed delay in cell growth of the deleted strain may be due to the increased cell death induced by H_2_O_2_, but also to an increase in cell cycle arrest. All of these experiments have confirmed the importance of Prn1 for cell survival and recovery during oxidative stress, such as mammalian Pirin (21) or *Streptomyces* PirA (17), corroborating a conserved function of this protein. Mammal Pirin is functionally similar to the oxidoreductase protein quercetin 2,3-dioxygenase, thus, this protein has been proposed to have an intrinsic oxidoreductase enzymatic activity (22). In depth *C. albicans* Prn1 sequence analysis using Pfam software identified, like mammal pirin, two conserved pirin domains (Fig. S5), which could indicate also a conserved intrinsic oxidoreductase activity supporting our described results.

### Prn1 absence increases ROS levels after H2O2 treatment

The DDA-MS proteomics approach for characterization of the proteome remodelling in *prn1Δ* and SN250 under oxidative stress showed a clear difference in the response of the two strains (Fig. 2 and Table 1). The GO term enrichment (Fig. 3 and Table S3) and the STRING analyses (Fig. 4) of the proteins that significantly increased in abundance for the two strains suggest a stronger oxidative stress response in the wild-type strain. Our hypothesis was functionally validated by the detection of higher intracellular ROS levels for the *prn1Δ* strain under oxidative stress (Fig. 6a) and a concomitant increase in apoptotic cells compared with the wild-type strain (Fig. 6b). Concordantly, mammal pirin has been also related to apoptosis programmed cell death (23). STRING software also detected clusters of mitochondrial oxidoreductase complex and heat shock proteins in the *prn1Δ* strain, which suggests that these proteins increase in abundance to counteract the absence of Prn1.

Proteins detected exclusively in the SN250 strain after H_2_O_2_ treatment with an opposite or non-significant change in the *prn1Δ* strain (Table 2 and Table 3) likely correlate with the presence of Prn1. A promising protein is Ald4 dehydrogenase, which is necessary for carnitine biosynthesis and proposed to be an important antioxidant in *S. cerevisiae* (24, 25). Another interesting protein was Rib2, which has pseudouridine synthase activity implicated in riboflavin biosynthesis. Riboflavin also has important antioxidant activity in both mammals (26) and yeasts (27). Ald4 and Rib2 increase in abundance in the SN250 strain after H_2_O_2_ treatment, suggesting that these antioxidants are trying to detoxicate the cell. In mammalian cells, Pirin is functionally similar to the oxidoreductase protein quercetin 2,3-dioxygenase. Quercetin is a flavonoid with antioxidant activity implicated in free intracellular ROS detoxification, thus, mammalian Pirin has been proposed to have intrinsic oxidoreductase enzymatic activity (22, 28). Prn1 may have conserved intrinsic antioxidant activity through riboflavin or carnitine flavonoids how mammalian Pirin acts on quercetin flavonoid.

In mammalian cells, the active conformation of Pirin (Fe^3+^) is crucial for NF-κB p65 transcription factor binding, which suggests that this protein may connect the oxidative stress response to the proteomic changes associated with this transcription factor (15, 16, 29). This could hint at the possible role of Prn1 as a translational coregulator involved in oxidative stress responses or apoptosis in *C. albicans*, similar to Pirin in mammalian cells. A possible relationship between NF-κB and the yeast retrograde response gene (RTG) signalling pathway has been proposed for *S. cerevisiae* (30). However, we did not identify any significant changes in the abundance of RTG pathway components in the wild-type strain, in which Prn1 is expressed (Table S6).

Surprisingly, the *PRN1* homologue genes have not shown a redundant function or expression. The proteomics analysis detected a slight but significant increase in Prn4 abundance (p-value) <0.05), which is the most similar to Prn1, in the wild-type strain after H_2_O_2_ treatment, but not in the *prn1Δ* after treatment, whereas Prn2 and Prn3 were not detected in any of the four conditions. Further studies will be necessary to discover their functions.

### Absence of Prn1 leads to an increase in proteasome activity and decrease in proteins related to translation after H_2_O_2_ treatment

Enrichment of protein catabolic process GO term proteins was observed in both strains due to the increase in free ROS after H_2_O_2_ treatment, which previous studies have described for the *C. albicans* SC5314 strain (12). In the SN250 strain, the most relevant proteins of this GO term were related to the transport of proteins from the Golgi to vacuole within the ubiquitin-dependent protein catabolic processes. However, for the *prn1Δ* strain, the most remarkably increased proteins were related to proteasome regulatory subunit (Pr26, Rpt4, and Rpn3), and some of these proteins were only detected in the treated *prn1Δ* strain (Fig. 3b and Table S1). Moreover, proteins that significantly increased in abundance only in the *prn1Δ* strain after treatment were enriched in protein catabolism GO term (Fig. S2). The higher proteasome activity in the mutant strain, possibly due to the higher ROS levels, was functionally validated (Fig. 6c), suggesting that increased proteasome activity is necessary to counteract the loss of Prn1 function. Proteins detected exclusively in the *prn1Δ* strain after treatment and with non-significant or an opposite change in the SN250 strain (Table 2 and Table 3) may be necessary to cell survival and recovery in the absence of Prn1. These proteins were related to protein catabolism and proteasome regulatory subunit. Also remarkable were two transcription factors (Mnl1 and Bas1) with functions that have not yet been completely described.

A decrease in proteins related to translation has been observed during the *C. albicans*-macrophage interaction (8, 31). In our experiment, we observed a clear decrease in the abundance of proteins related to both the translation GO term and nucleotide metabolic process GO term in the *prn1Δ* strain, but this was not accentuated in the SN250 strain (Fig. S1 and S3b). In the *prn1Δ* strain, proteins that were exclusively detected in basal conditions were related to ribosome biogenesis and RNA metabolic process (Table 2). The higher ribosome biosynthesis and protein translation in the wild-type strain, which was probably related to a faster cell recovery after stress, suggests that Prn1 could also be important in the response to host phagocytes.

### Differential expression of transcription factors between strains

Proteins detected only in one condition for each strain (Tables 2 and 3) revealed drastic differences in the abundance of several transcription factors (Mnl1, Nrg1, Bas1, Tif33, and orf19.1150). The mechanisms regulated by Tif33 or orf19.1150 have not been completely described. Bas1, which is implicated in filamentous growth and virulence, presented an inverse change in abundance between both strains, increasing in the *prn1Δ* mutant but not being detected in the wild-type strain in response to H_2_O_2_. The same pattern of expression was found for Mfg1, a regulator of filamentous growth involved in virulence. Mnl1 is involved in weak acid stress responses, regulating the expression of several genes, including Prn1 (32). In the *prn1Δ* strain, Mnl1 was only detected under oxidative stress, correlating with three Mnl1-regulated oxidative stress response proteins that significantly increased in abundance after treatment (Fig. S6a). Under oxidative stress, Nrg1, which is an Mnl1 antagonist, was not detected in the *prn1Δ* strain and, concordantly, 35 proteins repressed by Nrg1 presented a significant abundance increase. Among these proteins we identified six oxidoreductase activity proteins (Ali1, Gpd1, Gre2, Oye23, Sod1, and Yah1). Detailed lists of proteins regulated by Mnl1 or Nrg1 identified under each condition with their relative abundances are provided in Table S7. Mnl1 was only detected in the *prn1Δ* strain under oxidative stress, when Prn1 is absent, and was not detected in the treated SN250 strain, in which Prn1 is highly increased, whereas Nrg1 was only detected in the treated SN250 strain (Fig. S6b). We found nine Nrg1-repressed proteins that significantly increased in abundance in the treated *prn1Δ* strain, in which Nrg1 is absent, with respect to the treated SN250 strain (Table S7). In mammals, Pirin exerts its function in the oxidative stress response through Nrf2 modulation (14), but a *C. albicans* Nrf2 orthologue has not yet been described. Further experiments will determine whether Mnl1 could be the Nrf2 homologue.

### Prn1 may be related to mitochondrial oxidative stress detoxication under basal conditions

Comparative analysis of the proteomes of both strains under control conditions revealed that mitochondrial proteins with significantly changed abundance in the *prn1Δ* strain were related to the mitochondrial oxidative stress response (e.g., chaperones) and mitochondrial ribosome. In the SN250 strain, these proteins were involved in mitochondrial cell redox regulation and mitochondrial respiratory function (Fig. S3a). These results may indicate that the lack of Prn1 under basal conditions leads to increased mitochondrial oxidative stress due to the electron transport chain function. In *Streptomyces*, PirA has been identified as a negative regulator of the beta-oxidation pathway, decreasing oxidative stress (17). In eukaryotes, this process is produced in the inner mitochondrial membrane, so Prn1 could have the same function in *C. albicans*, regulating oxidative stress. Future studies will focus on determining the mitochondrial Prn1 function in the absence of oxidative stress.

The Mnl1 and orf19.1150 transcription factors also presented a significant difference in abundance between the two strains in the absence of H_2_O_2_. Another transcription factor, Swi3, a subunit of the SWI/SNF chromatin remodelling complex required for the expression of different genes in *S. cerevisiae* (33), significantly increased in abundance in the SN250 strain.

### *S. cerevisiae CUB1* orthologue orf19.4850 involvement in the oxidative stress response

In depth analysis of proteins expressed only in the SN250 strain after treatment (Table 2) highlighted *S. cerevisiae CUB1* orthologue orf19.4850, which was exclusively detected in this strain and this condition. In *S. cerevisiae*, this protein is involved in DNA repair and proteasome function (19). However, the molecular function, biological process and molecular component of orf19.4850 have remained unknown for *C. albicans*. Our experiments support a possible role of this protein in the oxidative stress response and conserved proteasome-related function (Fig. 7). Exclusive *S. cerevisiae* Cub1 orthologue detection when Prn1 is expressed suggest a possible relationship between them in the oxidative stress response.

### Prn1 is a relevant actor in the oxidative response

Taken together, these results reveal that Prn1 is a relevant component of the oxidative stress response in *C. albicans*. The function of Prn1 under basal conditions seems to be related to the regulation of mitochondrial redox. After H_2_O_2_ treatment, this protein significantly increases in abundance, correlating with increased ROS detoxification, which leads to greater cell survival and faster recovery after stress. In the absence of Prn1, proteasome activity is increased, translation-related proteins decrease in abundance, and different transcription factors change in abundance after treatment relative to the wild-type strain. In addition, we postulate a possible relationship between Prn1 and the *S. cerevisiae* orthologue Cub1 (Fig. 8).

**FIG 8.**
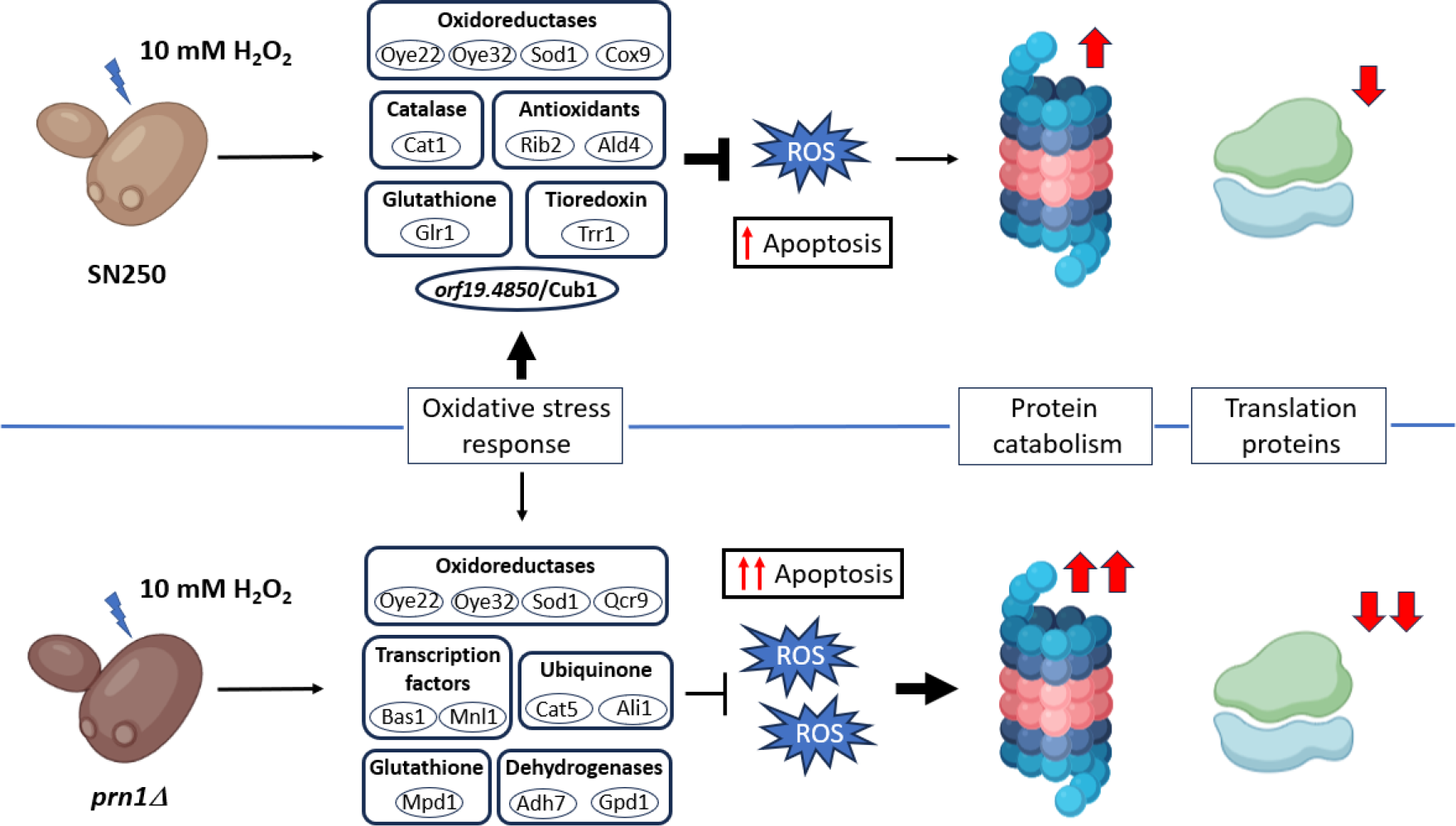
Schematic representation of the different responses of *C. albicans* to H_2_O_2_ in the presence or absence of Prn1. Wild-type strain presented an increased abundance of proteins from the three main detoxification systems (catalase, glutathione, and thioredoxin) in conjunction with different oxidoreductases and orf19.4850, a *S. cerevisiae CUB1* ortholog. The *prn1Δ* strain also had an increased abundance of most of oxidoreductases but not proteins related to other detoxification systems, such as Cat1 or Trr1. However, different dehydrogenases, ubiquinone biosynthesis proteins, and transcription factors also were increased in abundance. This different response causes increased ROS levels in the mutant strain, leading to increased cell death by apoptosis and necrosis and higher proteasome activity, together with putative decreased translation. The line thickness indicates the activation of each process. Adapted from (12).

## Methods

### Fungal strains and culture conditions

*Candida albicans* null *prn1Δ* mutant and the corresponding wild type strain (SN250) from Noble’s collection (18) were used for Prn1 function characterization assays. *Candida albicans* wild type SC5314 and *prn1Δ/PRN1* and *cub1Δ/CUB1* mutant strains were used for orf19.4850 function characterization assays. Cells were grown in conventional YPD rich medium (1% yeast extract, 2% peptone, and 2% glucose) at 30**°**C until reaching exponential phase (OD_600_= 0.8 – 1). Later, H_2_O_2_ was added at the appropriate times and concentrations, being incubated again at 30**°**C in rotatory shacking (180 rpm). H_2_O_2_ (Sigma-Aldrich) concentrations ranged from 5-7 mM for cell viability assays to 80 mM in drop growth assay. For growth curves, 10 μl of *C. albicans* culture (OD_600_= 0.04) were added to 180 μl of YPD medium and incubated for 72 h at 30**°**C with shacking. Regarding to drop growth assay, cells were treated with 80mM H_2_O_2_ and 4 μl of yeast culture were spotted every 3 min in YPD agar plates and incubated for 24h at 30**°**C.

### Viability assays

*C. albicans* cells were harvested after 200 min in the presence of 10 mM H_2_O_2_ and then 5 μl of propidium iodide (PI) (50 μg/ml) were added for 5 min at room temperature. Percentage of necrotic cells was measured by flow cytometry and results were analysed using FlowJo software. Statistical analysis was performed using a t-test.

### Cellular apoptosis

Cells were cultured and treated with 10 mM H_2_O_2_ for 50 min. Phosphatidylserine (PS) externalization was measured using annexin V-fluorescein isothiocyanate (FICT) in protoplast cells obtained applying standard techniques. Briefly, *C. albicans* cells were resuspended in a buffer containing 50 mM K_2_HPO_4_, 50mM dithiothreitol (DTT) and 5mM EDTA pH 7.2 at 30°C. Then, a buffer composed by 50 mM KH_2_PO_4_, 40 mM 2-mercaptoethanol, 0.15 mg/ml zymolyase 20T, 20 ml glusulase and 2.4 M sorbitol were added and incubated for 30 min. Protoplasts were stained following ApoAlert kit indications (TaKaRa) and the ratio of apoptotic cells was measured by flow cytometry. Results were analysed using FlowJo software. Statistical analysis was performed using a t-test.

### Cell disruption and protein extracts quantification

Control and treated cells were harvested and washed thrice in PBS. Next, lysis buffer (50 mM Tris-HCl [pH 7.5], 1 mM EDTA, 1 mM DTT, 150 mM NaCl, 10% protease inhibitors (Pierce) and 5 mM phenylmethylsulfonyl fluoride [PMSF]) was added to resuspend cell pellet. In the case of proteasome activity assay, no protease inhibitors or PMSF were added. Cell extracts were obtained by mechanical shaking in a Fast-Prep system (Bio101; Savant) 5 cycles of 30 s, applying glass beads (0.5 to 0.75 mm diameter). Samples were centrifuged for 15 min at 13,000 rpm to separate protein extracts from the cell debris and protein concentrations were measured using Bradford assay.

### Proteomic assay

*C. albicans* SN250 and *prn1Δ* strains were incubated for 200 min with 10 mM H_2_O_2_. 4 samples of both strains in each condition (control or treated) were obtained to perform the assay. After cell disruption, peptide digestion was performed using 50 μg of protein extracts (iST kit, PREOMICS) (34). In few words, samples were denaturalized, reduced and alkylated, later they were digested applying a trypsin/LysC mix and peptides were purified using a reversed-phase LC-MS column. Final peptide concentration on samples was quantified by fluorimetry using a Qubit4 system (Thermo Scientific). Mass spectra were acquired in a Q-Exactive HF hybrid quadrupole-Orbitrap mass spectrometer (Thermo Scientific) in a full-MS data-dependent acquisition (DDA) in a positive mode with Xcalibur 4.5 software. MS scans were acquired at m/z range of 350 to 1800 Da followed by data-dependent MS/MS scan (with a threshold of 0.01) of the 15 most abundance precursors with charges of 2-5 in MS scans for high-energy collision dissociation (HCD) fragmentation with a dynamic exclusion of 10 s and normalized collision energy (NCE) of 20. Peptide spectrum matches were filtered to a false-discovery rate (FDR) of >1%.

### Data analysis and GO enrichment analysis

Mass spectra (raw files) were processed using Proteome Discoverer 2.4 using the *Candida* Genome Database (CGD) (35) to generate a sample-specific peptide list. Mascot version 2.6 was used for the characterization and quantitative analysis of the yeast peptides. Search parameters included carbamidomethylation of cysteines as fixed modification, oxidation of methionine and N-terminal acetylation as variable modifications and a maximum of 2 missed cleavages allowed. Analysis for significant protein abundance changes between treated and their respective control samples were performed applying statistical qvalue <0.05. Determination of significant protein abundance changes between both treated samples were also performed applying statistical qvalue <0.05. Go enrichment analysis were carried out using GO Term Finder and GO Slim Mapper tools from Candida Genome Database (CGD) (36) Protein network clusters analysis was performed using STRING v.12.0 software.

### ROS detection

Yeast cells were incubated during 200 min with 10mM H_2_O_2_ and then washed thrice with cold PBS. Intracellular ROS were detected by staining treated cells with dihydrorhodamine 123 (DHR-123) (Sigma) at 5 μg/ml final concentration during 30 min. ROS species were detected by fluorescence microscopy counting >200 cells in each of the 3 biological replicates. Fiji – ImageJ software was utilized to quantify fluorescence signal taking account cell volume and subtracting background fluorescence of the image. Statistical analysis was performed using a t-test.

### Proteasome activity

SN250 and *prn1Δ* strains were treated with 10 mM of H_2_O_2_ during 200 min. Total extracts were obtained as previously described and later, proteasome chymotrypsin-like protease activity was measured in a fluorometric assay. Proteasome LLVY-R110 substrate (Proteasome 20S activity assay kit; Sigma-Aldrich) was added to 100 μg of total extracts during 2 h at 30°C, taking account manufacturer recommendations. Fluorescence signal was measured with BMG FLUOstar Galaxy (λex = 480 to 500 nm/λem = 520 to 530 nm). Statistical analysis was performed using a Mann-Whitney test.

## Data availability

The data set from proteomic analysis has been deposited in the ProteomeXchange Consortium via the PRIDE partner repository with the data set identifier PXD040804, project DOI: 10.6019/PXD040804 (37).

## Acknowledgements

This study was supported by grant PID2021-124062NB-100 from the Spanish Ministry of Science and Innovation. The proteomic analyses were performed in the Biological Techniques CAI, Proteomics Unit of the Complutense University of Madrid (UCM). The flow cytometry analyses were performed in the Biological Techniques CAI, Flow Cytometry and Fluorescence Microscopy Unit of the Complutense University of Madrid (UCM). V. Arribas acknowledges support by a contract from University of Salamanca through the European NextGenerationEU Fund and Spanish Ministry of Universities “Ayudas para la recualificación del sistema universitario español, modalidad Margarita Salas 2021-2022” and also to Microbiology and Parasitology Department of Complutense University of Madrid for the opportunity of a long-term stay.

## SUPPLEMENTAL FIGURES

**FIG S1.**
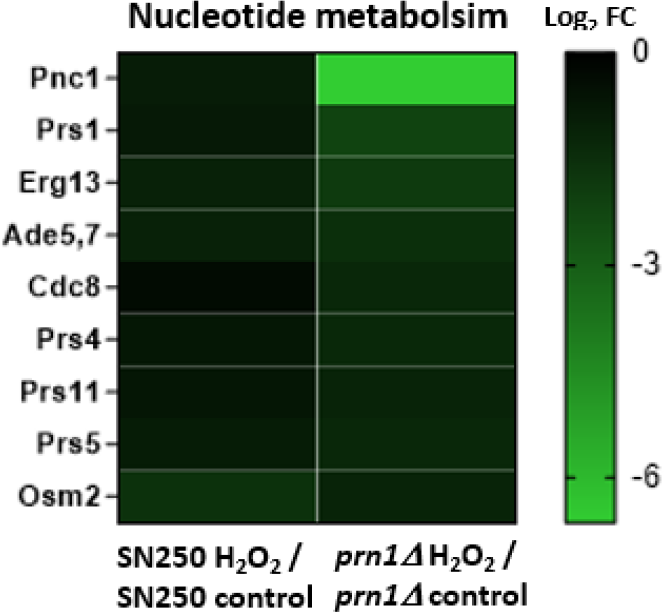
Hierarchical clustering heat maps of decreased nucleotide metabolism proteins after 200 min 10 mM H_2_O_2_ treatment in each strain.

**FIG S2.**
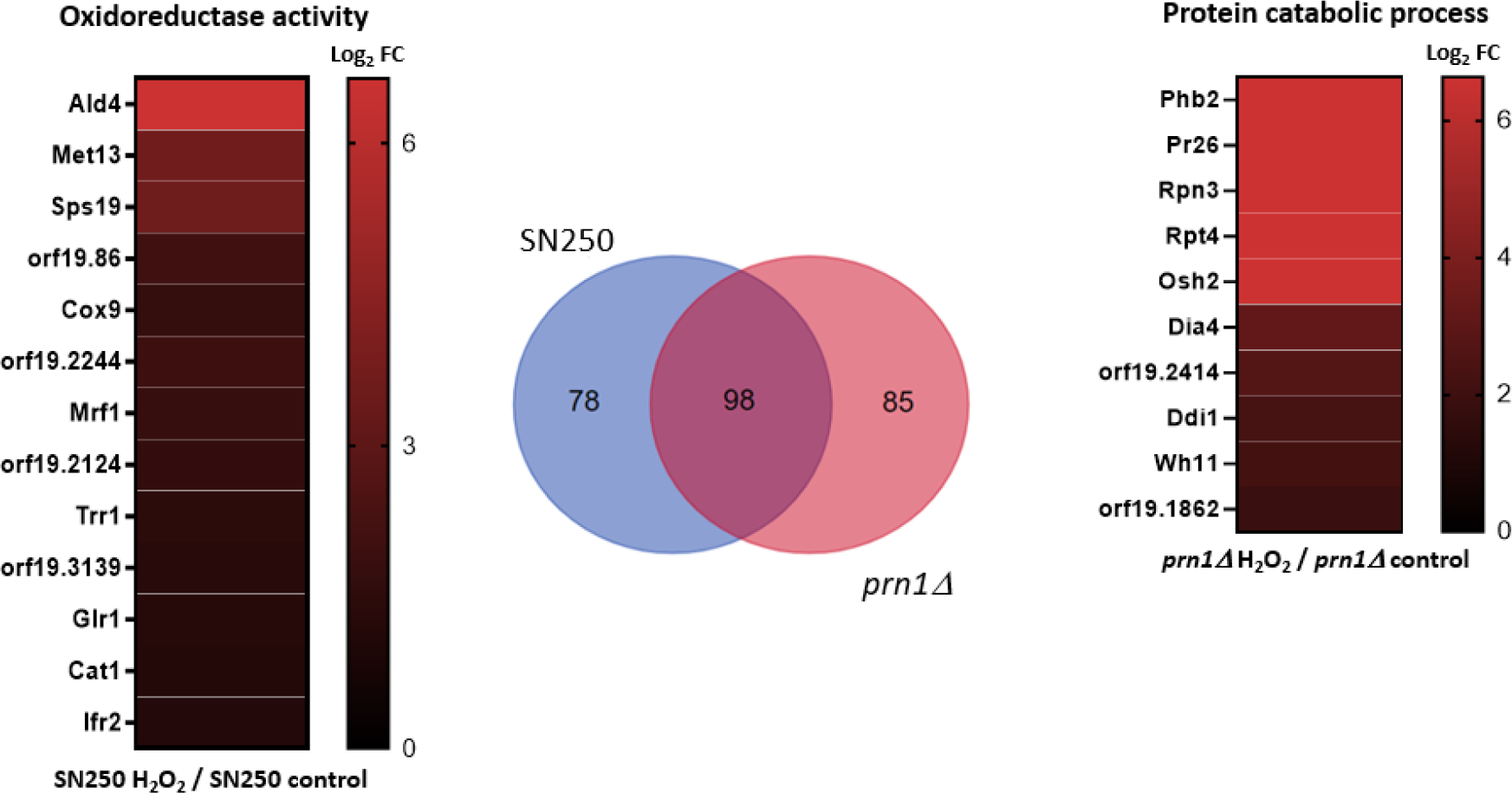
Hierarchical clustering heat maps of (a) increased oxidoreductase activity proteins in the SN250 strain and (b) increased protein catabolic process proteins in the *prn1Δ* strain after 200 min 10 mM H_2_O_2_ treatment.

**FIG S3.**
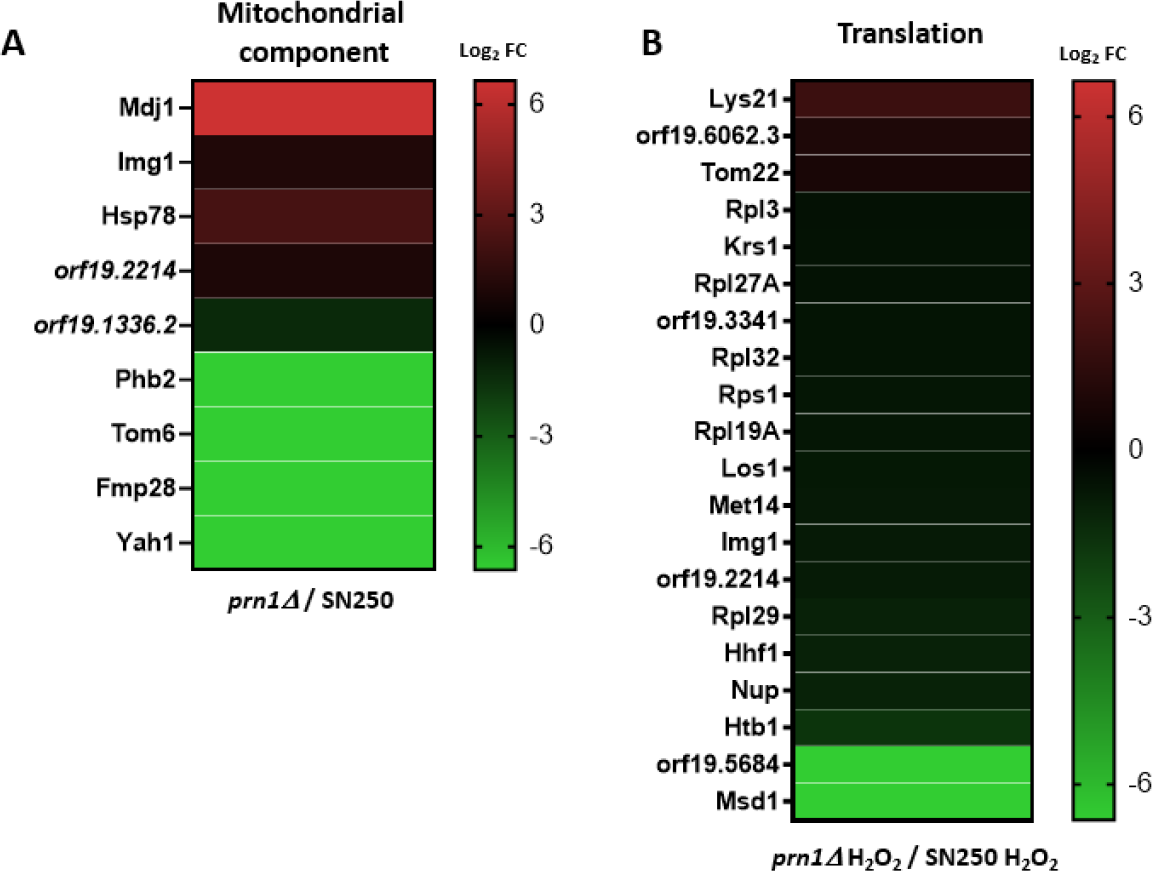
Hierarchical clustering heat maps of proteins that significantly changed in abundance grouped by Gene Ontology terms. (a) Basal conditions; (b) 10 mM H_2_O_2_ 200 min treatment.

**FIG S4.**
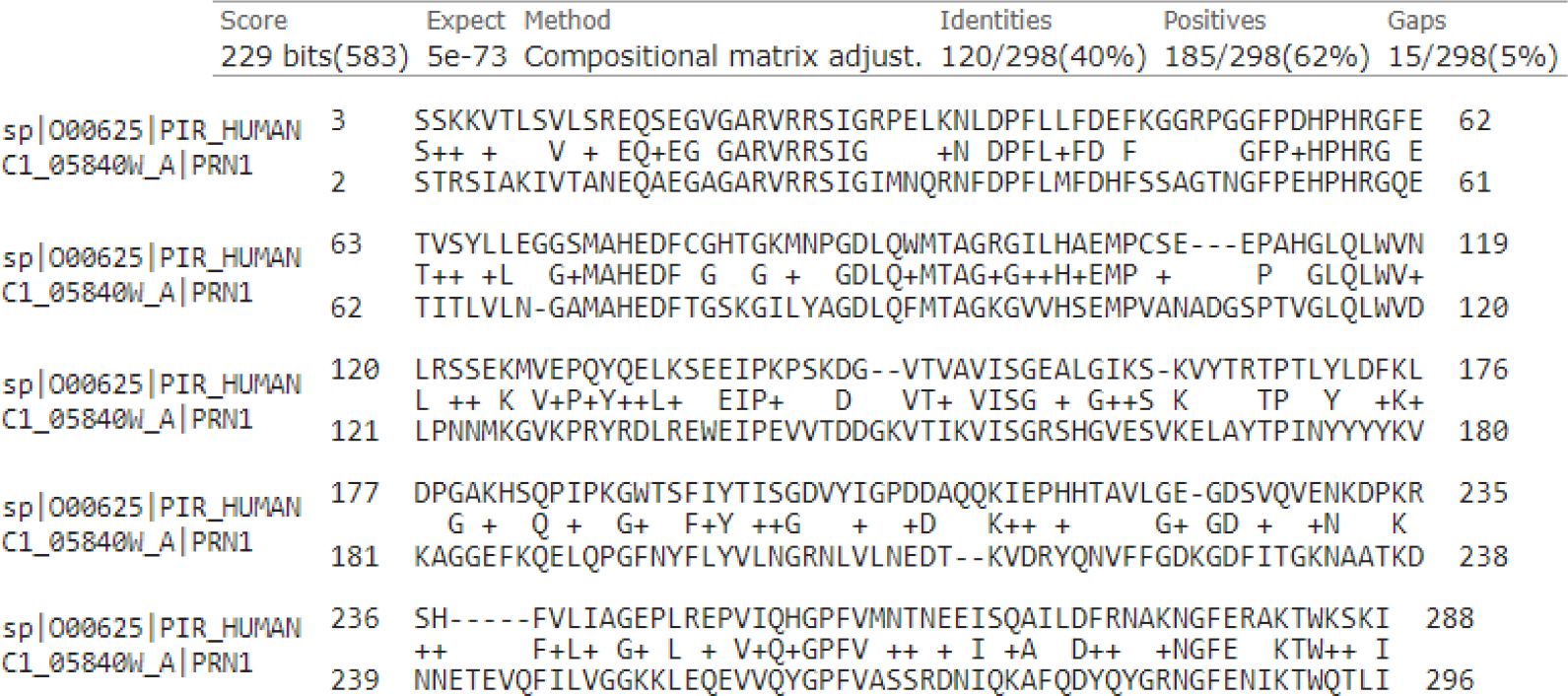
Human Pirin and *Candida albicans* Prn1 alignment using BLAST software. Prn1 presents 40% of identity with human Pirin

**FIG S5.**
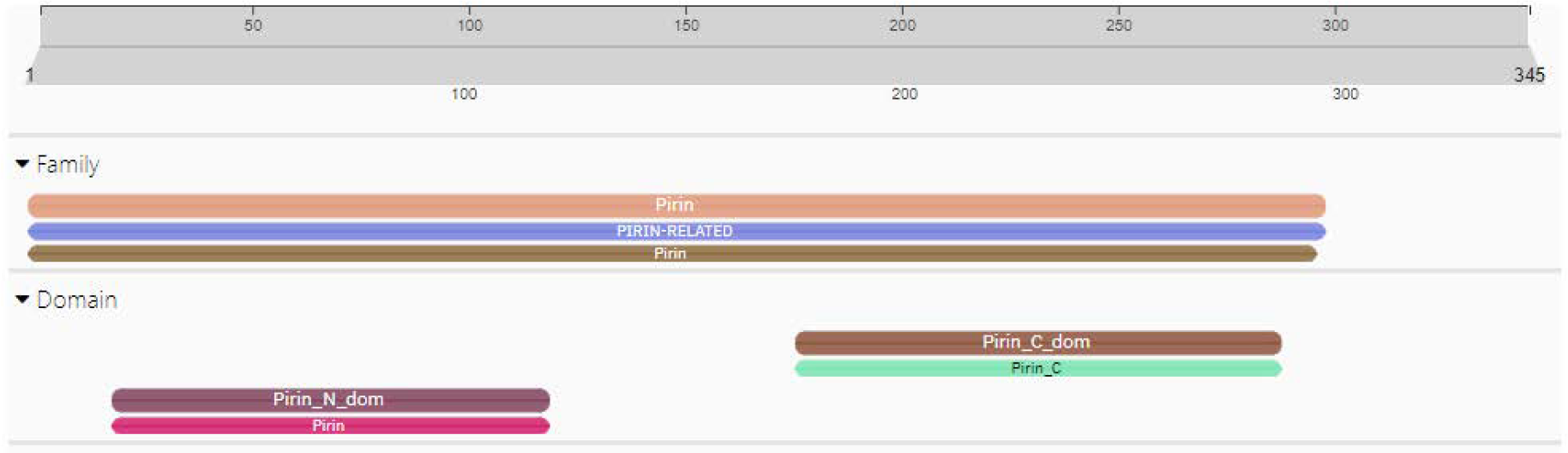
*Candida albicans* Prn1 sequence analysis using Pfam software. Prn1 presents one N-terminal pirin domain (amino acids 21-119) and another C-terminal pirin domain (amino acids 176-185).

**FIG S6.**
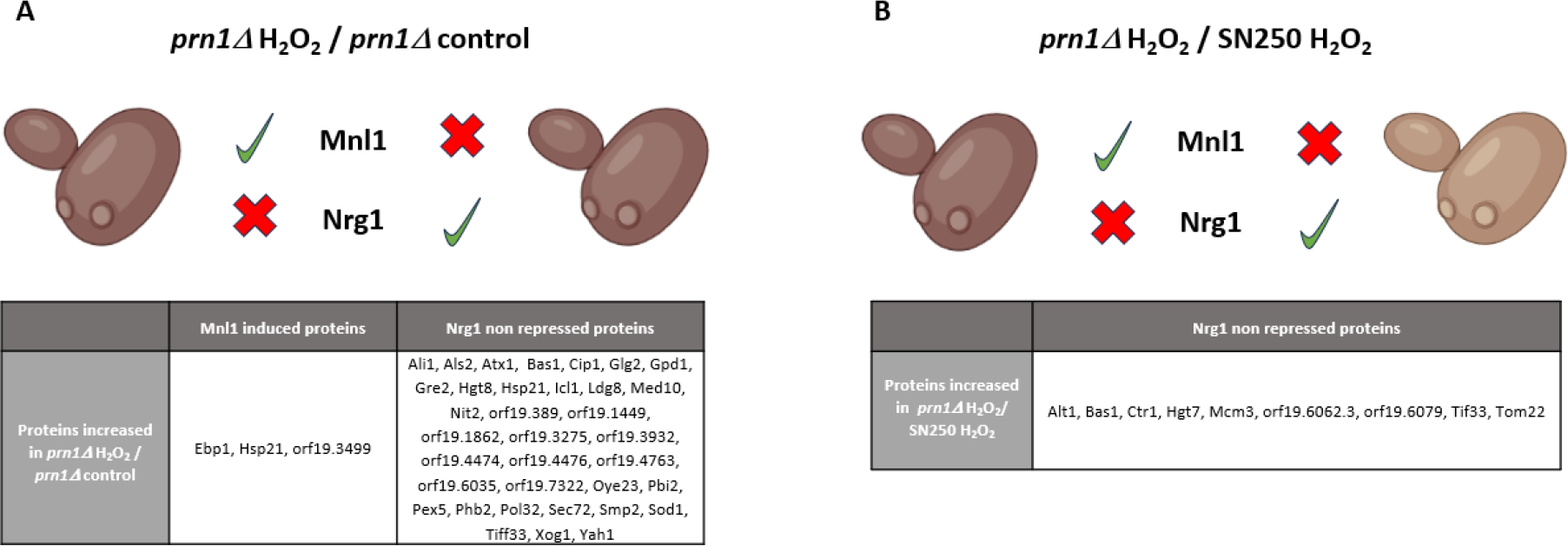
List of increased proteins regulated by Mnl1 and Nrg1 transcription factors in the *prn1Δ* strain treated with H_2_O_2_ with respect to (a) *prn1Δ* control and (b) SN250 H_2_O_2._ The relative ratio for each protein is presented in Table S1.

